# *MaUGT79* confers drought tolerance by regulating scopolin biosynthesis in plants

**DOI:** 10.1101/2023.11.20.567956

**Authors:** Zhen Duan, Fan Wu, Qi Yan, Shengsheng Wang, Yimeng Wang, Chris Stephen Jones, Pei Zhou, Caibin Zhang, Jiyu Zhang

## Abstract

The coumarin scopoletin and its glycosylated form scopolin constitute a vast class of natural products that are considered to be high-value compounds, distributed widely in the plant kingdom, they help plants adapt to environmental stresses. However, the underlying molecular mechanism of how scopolin is involved in the regulation of plant drought tolerance remains largely unexplored. Here, UDP-glycosyltransferase 79 (MaUGT79) was genetically mapped as the target gene by bulk segregant analysis sequencing (BSA-seq) from two *Melilotus albus* near-isogenic lines (NILs). MaUGT79 exhibits glucosyltransferase activity toward scopoletin. The expression of *MaUGT79* is induced by drought stress and it was found to mediate scopolin accumulation and reactive oxygen species (ROS) scavenging under drought stress. Moreover, the transcription of *MaUGT79* was demonstrated to be directly and positively regulated by MaMYB4, which is a key integrator of both scopolin biosynthesis and drought tolerance. Collectively, this study reveals that MaMYB4 is a positive regulator in drought stress by targeting the *MaUGT79* promoter and activating its expression to coordinately mediate scopolin biosynthesis and drought tolerance, providing insights into the regulatory mechanism for plant growth adaption to environmental changes through accumulation of scopolin.

## Introduction

In nature, plants are constantly exposed to various biotic threats and unfavorable growing conditions such as drought stress. To cope with drought stress, plants have evolved complex adaptive strategies. One such mechanism is the capacity to produce an impressive arsenal of stress-protective secondary metabolites (Sharma *et al*., 2019; Stringlis *et al*., 2019). Phenylpropanoids constitute a vast class of secondary metabolites that are considered to be high-value compounds due to their immense structural diversity and wide range of biological activities, and these compounds play important roles in the adaptation of plants to their environment (Doll *et al*., 2018; Liu, Xinyu *et al*., 2022). Coumarins, which are a rich source of medicines and therapeutic drugs, are important natural products of phenylpropane metabolism (Zhang *et al*., 2005), and are thought to be beneficial to plants by conferring resistance to herbivorous insects (Alonso *et al*., 2009) and pathogens (Perkowska *et al*., 2021), facilitating nutrient uptake (Tsai & Schmidt, 2017; Robe *et al*., 2021), shaping the composition of the root microbiome (Stringlis *et al*., 2019; Voges *et al*., 2019), and scavenging reactive oxygen species (ROS) (Doll *et al*., 2018). Coumarin also has a beneficial health effect on the body in optimal consumption (Jakovljević Kovač *et al*., 2021). However high concentrations of coumarin results in plants with poor palatability and dose-limiting toxicity (Liu *et al*., 2010), which may be a major limiting factor for the use of forage legumes. Even so, coumarin metabolism is the focus of global attention due to a growing array of applications for its products, which have broad pharmacological prospectives and are commonly used in traditional spices and flavoring agents (Huang *et al*., 2022). Consequently, there is a clear increasing trend in studies pertaining to the biosynthesis of coumarins (Tiwari *et al*., 2016).

The biosynthetic pathway of the coumarin-glycoside scopolin branches off from the phenylpropanoid biosynthetic pathway at the level of the hydroxycinnamoyl-CoAs and cinnamoyl-CoAs. The coumarin scopoletin is produced from C2-hydroxylation of the hydroxycinnamoyl-CoA esters of the side-chain through 2- oxoglutarate- dependent dioxygenase feruloyl-CoA 6’-hydroxylase 1 (F6’H1) activity (Kai *et al*., 2008) and then *trans*–*cis* isomerization and lactonization by the activity of COUMARIN SYNTHASE (COSY) (Vanholme *et al*., 2019). Finally, scopoletin can be glycosylated by UDP-glycosyltransferases (UGT) to form scopolin in the cytoplasm (Le Roy *et al*., 2016; Song *et al*., 2018).

Glycosylation provides the plant with an easy way to modify molecules, the process plays an important role in regulating the solubility, stability and biological activity of various small molecules and it is closely related to drought stress responses of plants (Bowles *et al*., 2005; Tognetti *et al*., 2010). Glycosylation reactions are mediated by UDP-glycosyltransferases (UGTs) that catalyze the transfer of an activated nucleotide sugar to acceptor aglycones to form glycosides (Song *et al*., 2018). Increasing evidence has indicated that *UGT* genes play pivotal roles in biosynthesis of phenolic compounds in many species (Dong *et al*., 2020; Adiji *et al*., 2021; Huang, X-X *et al*., 2021) particularly in response to abiotic stresses (Rehman *et al*., 2018). However, only a few coumarins and their corresponding UGTs have been functionally characterized due to the presence of hundreds of UGT-encoding genes in most plant species and the substrate promiscuity of UGT enzymes (Sun *et al*., 2019; Krishnamurthy *et al*., 2020). *Nicotiana tabacum* UGT73A1 and UGT73A2 have substrate specificity toward coumarins forming scopolin and esculin (Li *et al*., 2001). *Tobacco-O-glucosyltransferase* (*TOGT*)-mediated glucosylation is required for scopoletin accumulation in cells surrounding tobacco mosaic virus (TMV) lesions, where this compound could both exert a direct antiviral effect and participate in reactive oxygen intermediate buffering (Chong *et al*., 2002). The glycosylation activity of UGT73C7 results in the redirection of phenylpropanoid metabolic flux to the biosynthesis of hydroxycinnamic acids and coumarins, promoting *SNC1*-dependent arabidopsis immunity (Huang, X-X *et al*., 2021). Mounting evidence suggests that *UGT* is the key conduit for regulating coumarin biosynthesis in plants.

MYB transcription factors (TFs) belong to one of the largest and most important gene families, which regulate development under changing environmental conditions, primary and secondary metabolism, and plants response to stresses. Several members of the R2R3 MYB family have been reported to be involved in the biosynthesis of phenylalanine and phenylpropanoid-derived compounds (Chen *et al*., 2019). MYB4 is reported to negatively regulate itself by binding to its own promoter (Zhao *et al*., 2007). AtMYB7, a homolog of AtMYB4, repressed the expression of *UGT* genes that encode key enzymes in the flavonoid pathway (Fornale *et al*., 2014). MYB72 regulates the biosynthesis of iron-mobilizing phenolic compounds, after which BGLU42 activity is required for their excretion into the rhizosphere (Stringlis *et al*., 2018). One of the very few works dealing with the regulation of scopolin production by transcription factors is about the antagonizing transcription factors MYB12, promoting flavonol synthesis, and MYB4, suppressing flavonol synthesis and thus promoting scopoletin production (Schenke *et al*., 2011). However, the functions of MYB TFs in scopolin biosynthesis remains largely unknown, and the associated regulatory mechanisms is still a mystery.

*Melilotus albus* is a diploid species with 8 chromosomes (2*n*=16) and a sequenced genome of ∼1.05 Gb (Wu *et al*., 2022). It is an excellent rotation crop because it has been used for both forage production and soil improvement (Zabala *et al*., 2018), and it is drought tolerant, winter hardy and grows in practically all soil types (Kulinich, 2020). Coumarin content varies significantly among different *Melilotus* species (Nair *et al*., 2010), and ranges from 0.2% to 1.3% of the dry matter within *M. albus* (Zhang *et al*., 2018). Recent studies have suggested that coumarins play a crucial role in plant drought tolerance (Rangani *et al*., 2020); however, the underlying molecular mechanisms are still largely unknown. For *M. albus*, most studies were about the conventional breeding of low coumarin content germplasm (Luo *et al*., 2018; Zhang *et al*., 2018) rather than deciphering the molecular mechanism. In our previous studies, differentially expressed unigenes and miRNAs involved in coumarin biosynthesis have been identified in *M. albus* (Luo *et al*., 2017; Wu *et al*., 2018). The *UGT* gene family of *M. albus* has been identified (Duan *et al*., 2021). The key enzymes in the coumarin biosynthesis pathway have been identified in *M. albus* near-isogenic lines (NILs), JiMa46 and JiMa49 (Wu *et al*., 2022). However, there has been little progress on functional analysis of coumarin biosynthesis genes. Therefore, filling the knowledge gap in molecular mechanisms of coumarin biosynthesis is particularly important.

In this study, we identified a UDP-glycosyltransferase encoding gene which was previously uncharacterised in *M. albus*, *MaUGT79*. We investigated the molecular functions of the *MaUGT79* gene in scopolin biosynthesis and drought tolerance through the generation and characterization of transgenic hairy roots over-expressing *MaUGT79*, as well as RNA interference (RNAi)-mediated knockdown of *MaUGT79*. We also found that MaMYB4 positively regulates the plants drought stress response through coordinately activating *MaUGT79*-involved scopolin biosynthesis. Our results unravel the mechanism of *MaUGT79*-mediated scopolin biosynthesis and drought tolerance, provide new genetic resources for enhancing drought tolerance in *M. albus* breeding and potentially contribute to sustainable agriculture in terms of weathering drought stresses.

## Results

### An unknown UDP-glucosyltransferase, MaUGT, was genetically mapped based on the BSA-seq in *M. albus*

Scopolin contents in different tissues for two NILs (JiMa46 and JiMa49) of *M. albus* were analyzed by HPLC. Significantly higher levels of scopolin in leaf, stem and root were observed in JiMa49 than in JiMa46 (Figure 1A). To identify genes involved in the scopolin biosynthesis of *M. albus*, the BSA-seq was carried out. We found with high confidence that between JiMa46 and JiMa49, major polymorphic loci (SNPs and InDels) mapped at the 67,539,223 to 67,540,565 positions on chromosome 5, a region which was annotated as the UDP-glucosyltransferase (UGT) encoding gene *Malbus0502448.1*. This gene, with an unknown function, was assigned as the candidate gene underlying these loci (Figure 1B). We obtained the nomenclature-appropriate names of 189 full-length *UGT* genes based on chromosomal position from the *M. albus* genome (Duan et al., 2021), *Malbus0502448.1* is designated as *MaUGT79*. To further confirm the gene, the CDS of *MaUGT79* was amplified from JiMa46 and JiMa49 and the structures were analyzed. Sequence comparison among the UDP-glucosyltransferase encoding genes identified many nucleotide variations and InDels between the two NILs, which also exhibited frameshift mutations in the gene. *MaUGT79*_JiMa46 shows a ten-base deletion at the N-terminal region compared with *MaUGT79*_JiMa49, which causes the early termination of *MaUGT79*_JiMa46 translation at the 121 amino acid position and the loss of the conserved Plant Secondary Product Glycosyltransferase (PSPG) box domain, while the full translation of *MaUGT79*_JiMa49 can proceed, generating a 463 amino acid peptide (Figure 1C). Therefore, we assumed that the InDel which induced the premature termination of translation may affect protein function and thereby cause a loss of the *MaUGT79* gene function for glucosyltransferase in JiMa46. Hence, we considered that *MaUGT79* was the most likely candidate gene responsible for scopolin biosynthesis.

**Figure 1.**
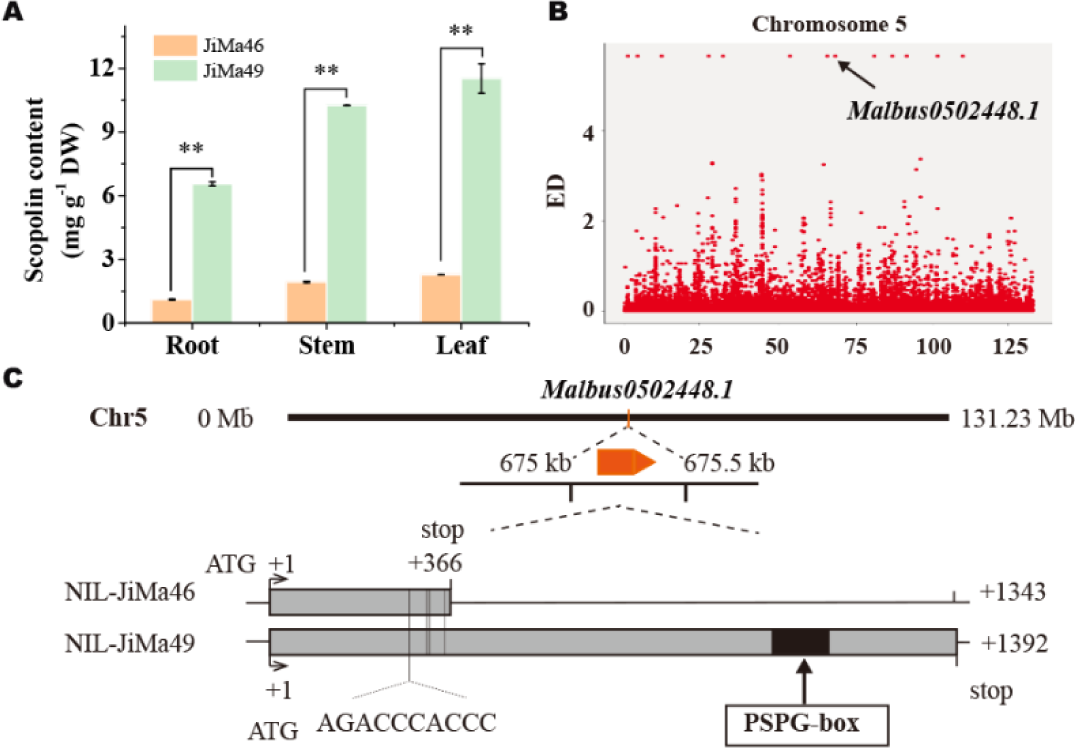
Discovery of a specific scopoletin UDP-glycosyltransferase in *M. albus*. A) Scopolin content in root, stem and leaf tissues of 2-month-old plants at the flowering stage of the two NILs, JiMa46 and JiMa49. The error bars indicate the SD values from at least three repetitions. Significant differences were detected by Student’s t-test: **, *P*<0.01. B) Identification and localization of the scopolin biosynthesis locus between the two NILs based on BSA. The annotated gene was identified with SNPs and InDels between NILs JiMa46 and JiMa49. Each point represents an individual SNP locus. C) Physical position, gene structure, polymorphisms between JiMa46 and JiMa49 of *Malbus0502448.1*. Mutational changes in JiMa49 are indicated. Exon and the highly conserved plant secondary product glycosyltransferases (PSPG) box of plant glycosyltransferases are indicated in grey and black boxes, respectively. Nucleotide polymorphisms are indicated at their corresponding positions in the coding sequences.

*MaUGT79* contains a 1392 bp open reading frame (ORF) encoding a protein with a molecular mass of 51.86 kDa and a pI of 5.60. To further predict the function of *MaUGT79*, we aligned its protein sequence with similar previously characterized UGTs and carried out a phylogenetic analysis. Members of the UGT enzyme family consist of two similar (N- and C-terminal) domains, each possessing several alpha helices and beta strands (Adiji *et al*., 2021). Multi-sequence alignment comparison analysis revealed that MaUGT79 had high amino acid identity within the PSPG box ‘WAPQ- 2x-IL-x-H-5x-F-2x-HCGWNS-x-LE-4x-G-4x-TWP-4x-Q’ near the C-terminal end (Figure 2A), which binds with the UDP portion of the sugar donor during catalysis. At the N-terminal region, MaUGT79 possesses a critical catalytic His19 that is universally conserved across all plant UGTs. A neighboring residue, Pro22, was also identified that is either crucial for interacting with the acceptor substrate during catalysis or can partly define donor substrate acceptability (Shao *et al*., 2005; Osmani *et al*., 2008) (Figure 2A). In addition, we carried out phylogenetic analyses based on the amino acid sequences of MaUGT79 and a set of UGTs that had been systematically screened for activity with a variety of hydroxylated benzoic acids, including coumarins (Song *et al*., 2016). Phylogenetic analysis indicated that MaUGT79 clustered in the same branch as AtUGT89B1 and AtUGT89A2, belonging to group B, which are closely related to group D that consists of known coumarin UGTs. UGT73B3 and UGT73B4 have been predicted as hydroxybenzoate glycosyltransferases, they readily glucosylate coumarins like their phylogenetic neighbors FaGT7 (Griesser *et al*., 2008) and TOGT1 (Langlois-Meurinne *et al*., 2005) (Figure 2B). Thus, the localization of MaUGT79 within this cluster would be consistent with a role in glycosylation of coumarin or other hydroxybenzoates.

**Figure 2.**
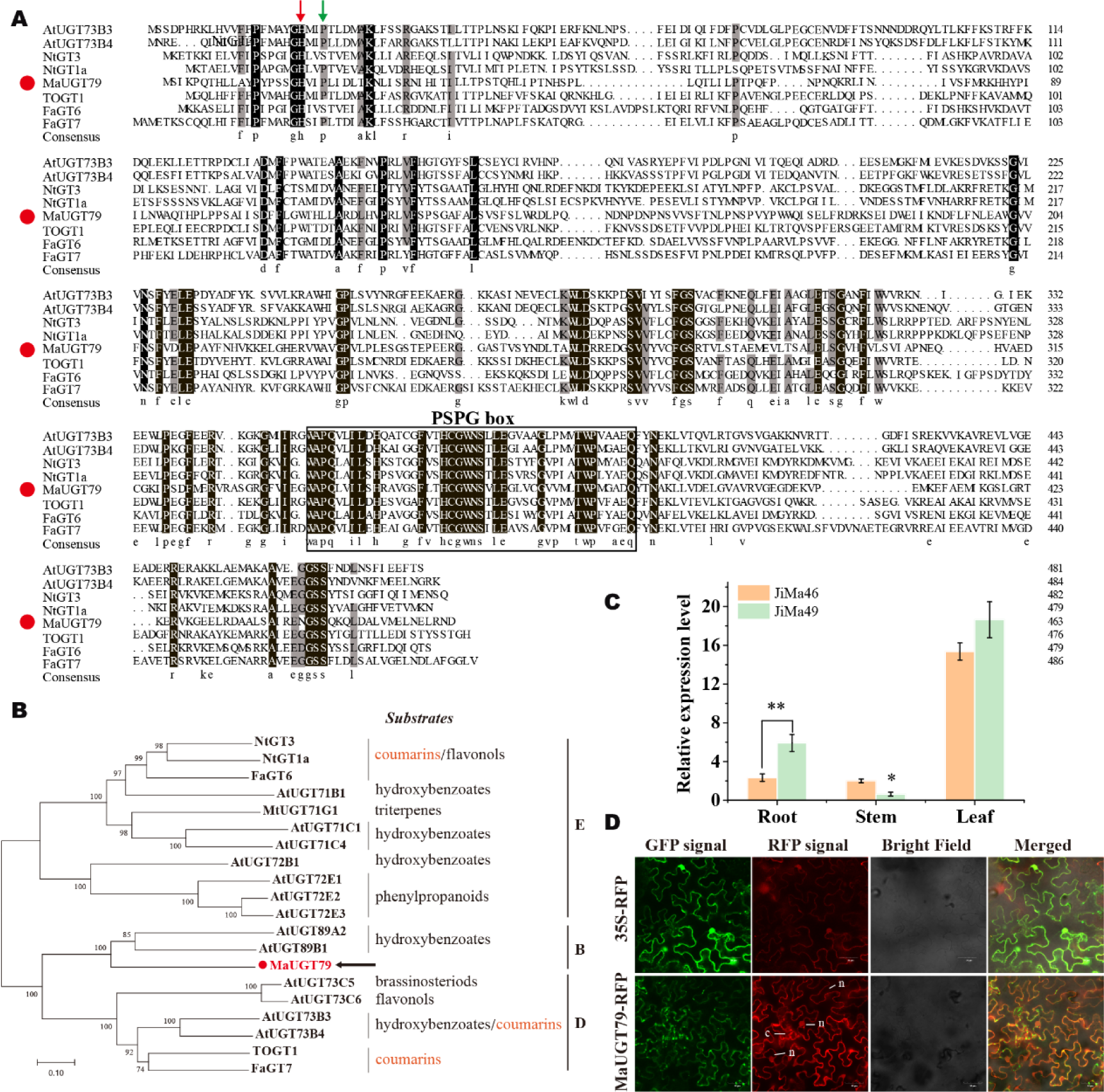
Amino acid sequence alignment, phylogenetic analysis, subcellular localization of MaUGT79. A) Amino acid alignments of MaUGT79 with other identified UDP-glycosyltransferase (UGT) proteins involved in coumarin biosynthesis including AtUGT73B3, AtUGT73B4, NtGT3, NtGT1a, TOGT1, FaGT6 and FaGT7. The red arrow at the N-terminus indicates the presence of a critical catalytic His that is universally conserved across all plant UGTs. A neighboring residue, believed to be important in contributing to sugar donor recognition, is indicated by the green arrow. The black rectangle at the C-terminus indicates the domains of the PSPG box. Numbers indicate the last residue in each line. Identical residues are highlighted by a black background and similar residues are highlighted by a grey background. The red dots indicate MaUGT79. B) Phylogenetic tree of MaUGT79 together with other functionally characterized UGTs. The tree is constructed using the Neighbor–Joining method by the MEGA W software. Numbers indicate bootstrap values for 1000 replicates. The GenBank accession numbers of the UGT proteins are: AtUGT73B3 (AAL32831); AtUGT73B4 (BAE99671); NtGT3 (BAB88934); NtGT1a (BAB60720); TOGT1 (AAK28303); FaGT6 (ABB92748); FaGT7 (ABB92749); AtUGT71B1 (BAB02837); MtUGT71G1 (RHN56458); AtUGT71C1 (AAC35226); AtUGT71C4 (AAG18592); AtUGT72B1 (AAK25972); AtUGT72E1 (AAK83619); AtUGT72E2 (BAA97275); AtUGT72E3 (AAC26233); AtUGT89A2 (CAB83309); AtUGT89B1 (AEE35520); AtUGT73C5 (AAD20156); AtUGT73C6 (AAD20155). The red dot indicates MaUGT79. C) Relative expression levels of *MaUGT79* in root, stem and leaf tissues of 2-month-old plants at the flowering stage of the two NILs, JiMa46 and JiMa49. Data are normalized by β*-tubulin*. Data are shown as the mean (n = 3), and significant differences were detected by Student’s t-test: *, *P*<0.05 or **, *P*<0.01. (d) The subcellular localization of MaUGT79 in *N. benthamiana* leaves. *MaUGT79* was fused to RFP. The fluorescence was observed under a confocal laser scanning microscope. Scale bars indicate 50 µm.

### Cytoplasm and nuclei localized *MaUGT79* is highly expressed in *M. albus* leaf and root tissues

The relative expression levels of *MaUGT79* in root, stem and leaf tissues between the genotypes JiMa46 and JiMa49 were analyzed (Figure 2C). The highest expression level was detected in leaf and root of JiMa49, where expression was significantly greater than in JiMa46, suggesting that *MaUGT79* may function in different tissues for the glycosylation process. We also examined the subcellular location of *MaUGT79* and found that the RFP signal was predominantly localized in the cytoplasm and nucleus (Figure 2D), suggesting a role for *MaUGT79* in scopolin metabolism in the cytoplasm. Certainly, the possibility of the partial diffusion of this protein back to the nuclei cannot be excluded.

### MaUGT79 exhibits scopoletin glucosyltransferase activity

Previous studies have indicated that the amino acids in the N-terminal region of UGTs are crucial for interacting with the sugar acceptor substrate and that the PSPG box in the C-terminal region of UGTs plays a key role in sugar-donor binding (Osmani *et al*., 2009). MaUGT79 is a member of the UDP-dependent glycosyltransferase family, which mainly uses UDP-glucose as a sugar donor to catalyze the glucosylation of plant secondary metabolites (Gachon *et al*., 2005; Bowles *et al*., 2006). We thus heterologously expressed MaUGT79 in *E. coli* and purified the protein (Figure S1), and then conducted a substrate feeding assay using *o*-coumaric acid, esculetin, umbelliferone and scopoletin as the possible substrate, and UDP-glucose as a sugar donor to explore the ability of the recombinant protein to glucosylate coumarins. HPLC analysis of reaction products showed no new peaks were produced when using *o*- coumaric acid and umbelliferone as substrates, indicating that MaUGT79 could not catalyze the glycosylation of *o*-coumaric acid and umbelliferone. When using esculetin as a substrate, a small new peak was generated in the reaction products, suggesting that MaUGT79 had a weak catalytic capacity for esculetin. When scopoletin was used as a substrate, scopolin was produced with exactly the same retention time of 13 min as the scopolin authentic standard (Figure 3), indicating that MaUGT79 mainly had glycosylation activity against scopoletin and could catalyze the conversion of scopoletin to scopolin.

**Figure 3.**
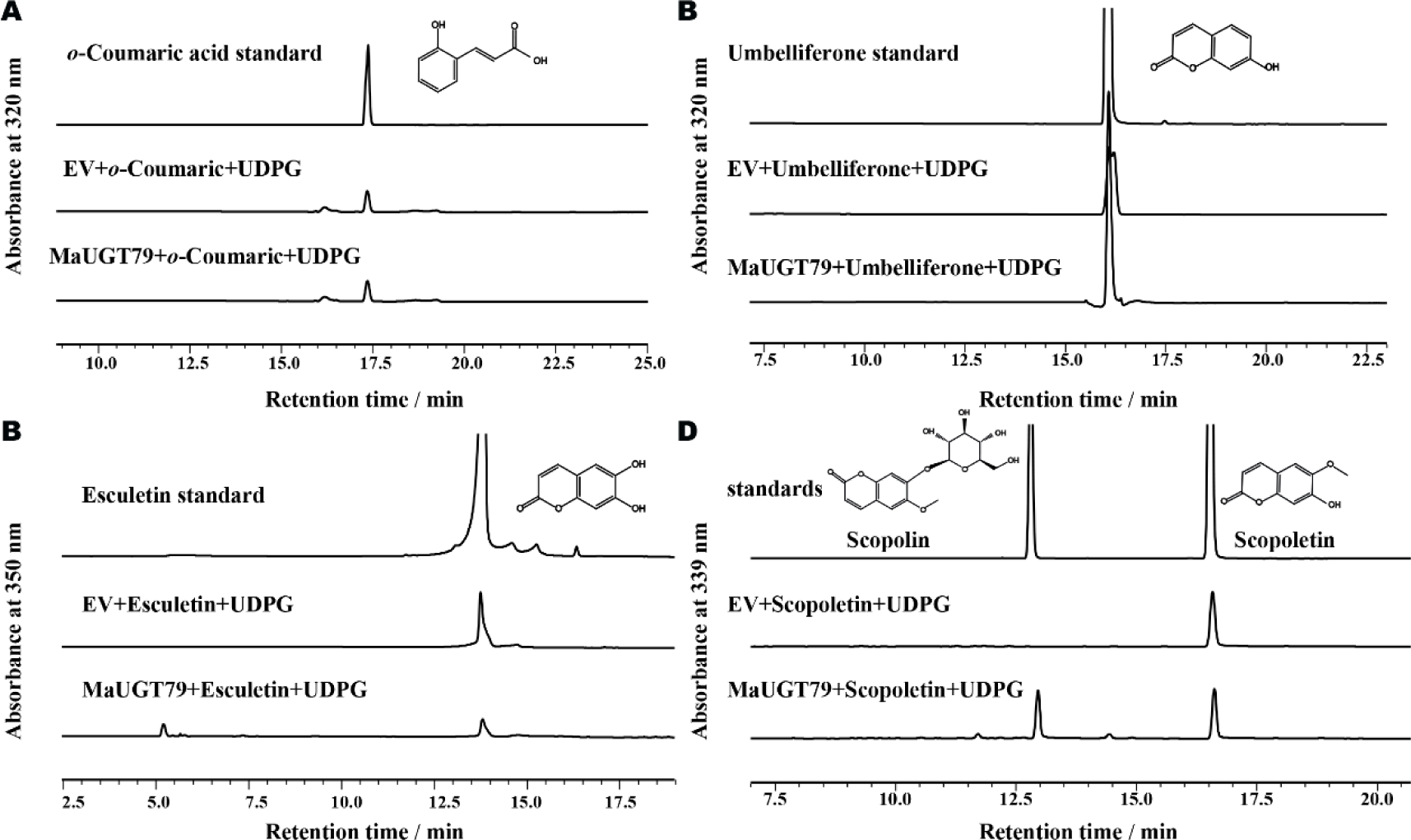
The *in vitro* glucosylating activity of MaUGT79 toward different coumarins. HPLC analyses of the reaction products catalyzed by fusion protein MaUGT79 with *o*- coumaric acid A), umbelliferone B), esculetin C), scopoletin D) compared with authentic standards. UDP-glucose was used as the sugar donor. Empty vector was used as a negative control. The authentic *o*-coumaric acid, umbelliferone, esculetin, scopolin and scopoletin were used as the standards.

### *MaUGT79* silencing decreased scopolin accumulation, while its overexpression enhanced scopolin biosynthesis

Here, *A. rhizogenes*-mediated hairy root transformation was successfully developed in *M. albus* to further investigate whether *MaUGT79* contributes to scopoletin glucosylation *in vivo*. The transgenic hairy roots were identified by genomic PCR, RFP signal detection, and qRT-PCR (Figure S2, Figure 4A). Transgenic hairy root OE- *MaUGT79* lines with relatively high expression and RNAi lines with low expression were selected for functional characterization, while three independent *M. albus* hairy root lines that express the empty vector were used as a control (EV). The total endogenous scopolin from the OE-*MaUGT79* lines and RNAi lines was extracted and analyzed by HPLC. We found that the scopolin accumulation in OE-*MaUGT79* lines is much higher than that of EV lines. On the other hand, the scopolin level in RNAi lines was almost half that detected in EV lines (Figure 4B). These data indicate that MaUGT79 catalyzes scopoletin glucosylation in *M. albus*. Then we assayed the expression levels of genes related to the phenylpropanoid pathway. As shown in Figure 4C, the expression levels of genes involved in the central phenylpropanoid pathway (*4CL1*, *CCoAOMT1*), and scopoletin biosynthesis (*F6’H1*, *BGLU42*) were obviously higher in OE-*MaUGT79* lines and reduced in RNAi lines compared with those in EV lines. We conclude that *MaUGT79* is indispensable for scopolin biosynthesis.

**Figure 4.**
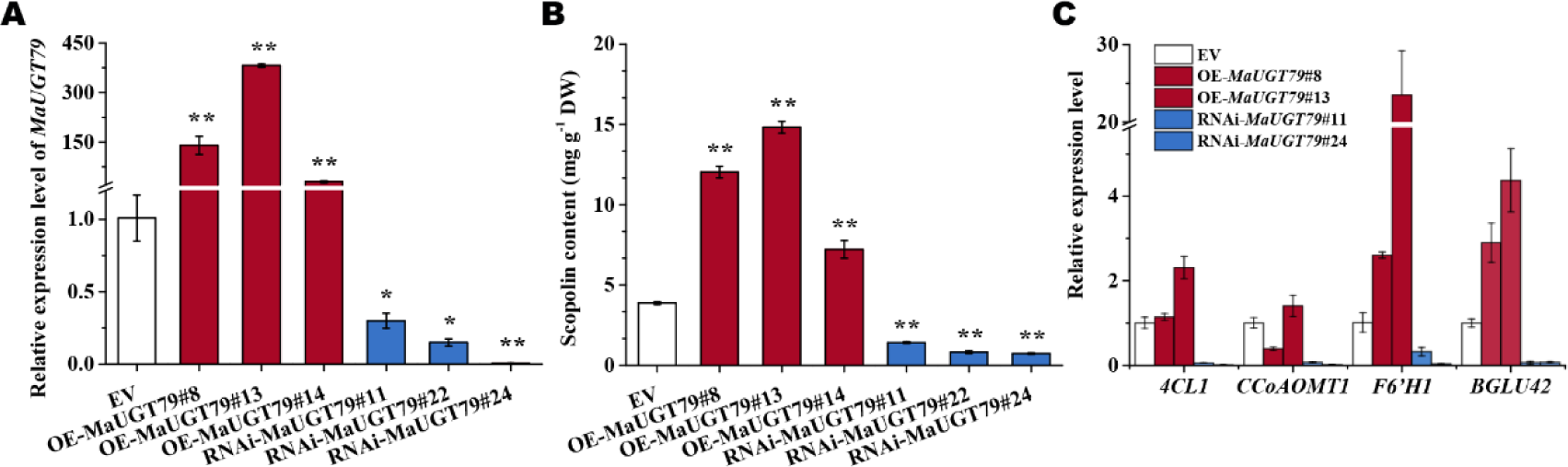
Over- or knockdown expression of *MaUGT79* alters scopolin content in *M. albus* hairy roots. A) Analysis of *MaUGT79* expression levels and B) scopolin content in control (EV), overexpression (OE) and RNAi transgenic hairy roots. C) Gene expression levels of four scopolin biosynthesis genes (*4CL1*, *CCoAOMT1*, *F6’H1*, and *BGLU42*) in 21-d-old hairy roots. qRT-PCR was performed to detect gene expression levels. Data are normalized by β*-tubulin*. Data are shown as the mean (n=3). The error bars indicate the SD values from at least three repetitions. Significant differences were detected by Student’s t-test: *, *P* < 0.05 or **, *P* < 0.01.

### *MaUGT79* contributes to drought stress tolerance through modulating scopolin biosynthesis

Coumarins are among the most bioactive plant secondary metabolites that serve as well-known antioxidants and show a response to drought stress by protecting plants against oxidative damage (Qin *et al*., 2019; Rangani *et al*., 2020; Patel *et al*., 2021). Applied exogenous scopolin increased scopoletin and scopolin content (Figure S3A) and decreased the MDA and O_2-_ content under 30% PEG6000 treatment and drought stress in EV and RNAi-*MaUGT79* hairy roots (Figure S3), indicating a positive role of scopolin reducing oxidative damage and promoting ROS scavenging under drought stress. Given that *MaUGT79* plays a significant role in modulating scopolin profiles, we asked whether *MaUGT79* is necessary for drought stress tolerance. First, an analysis of the expression of *MaUGT79* under drought stress was performed. We found that the expression of *MaUGT79* in *M. albus* was highly induced by 30% PEG treatment (Figure S4), and the conferring of drought tolerance by *MaUGT79* was confirmed in a yeast system (Figure 5A). To investigate how *MaUGT79* and its glycosylated scopolin affect drought stress tolerance, transgenic hairy root OE-*MaUGT79* lines with relatively high expression and RNAi lines with low expression levels were selected for functional characterization, while transgenic hairy roots containing an empty vector were used as a control. No phenotypic differences were observed between the plants with OE-*MaUGT79* and RNAi-*MaUGT79* hairy roots and control plants under normal growth conditions (the relative water content was 82.13%). After 3 days of 30% PEG6000 treatment and 22 days for water-deficit treatment (the relative water content was 25.13%), the leaves of control plants displayed slight wilting and necrosis, the leaves of the plants with RNAi-*MaUGT79* transgenic hairy roots withered and were yellow, while no obvious damage was observed in the plants with *MaUGT79*- overexpressing transgenic hairy roots (Figure 5B, H). The survival rates of the plants with *MaUGT79*-overexpressing transgenic hairy roots were 75-86.67%, whereas only 7.01% of RNAi-*MaUGT79* transgenic hairy roots plants survived. Control plants had a 52.22% survival rate 22 days after drought stress induction (Figure 5J). Nitroblue tetrazolium (NBT) staining indicated that the control hairy roots displayed more severe damage in comparison with OE-*MaUGT79* hairy roots (Figure 5C), which was consistent with the result of O_2-_ content (Figure 5e, n). MDA and H_2_O_2_ contents in OE- *MaUGT79* transgenic hairy roots decreased significantly relative to control hairy roots under 30% PEG6000 treatment and drought stress, while they increased in RNAi- *MaUGT79* transgenic hairy roots (Figure 5D, F, L, M). *MaUGT79*-overexpression lines treated with 30% PEG6000 and drought stress showed an increase in scopolin content and RNAi-*MaUGT79* lines showed a decreased scopolin content (Figure 5G, L). Therefore, we conclude that the reduced scopolin content in RNAi-*MaUGT79* hairy roots weakens the ROS scavenging activity and subsequently reduces drought tolerance. Consistently, drought marker genes, such as *MaCOR47*, *MaRD29A*, *MaLEA3*, *MaP5CS1*, *MaRD29B* and *MaDREB2B*, exhibited significantly elevated expression levels in two OE-*MaUGT79* lines upon exposure to 30% PEG6000 treatment (Figure S5).

**Figure 5.**
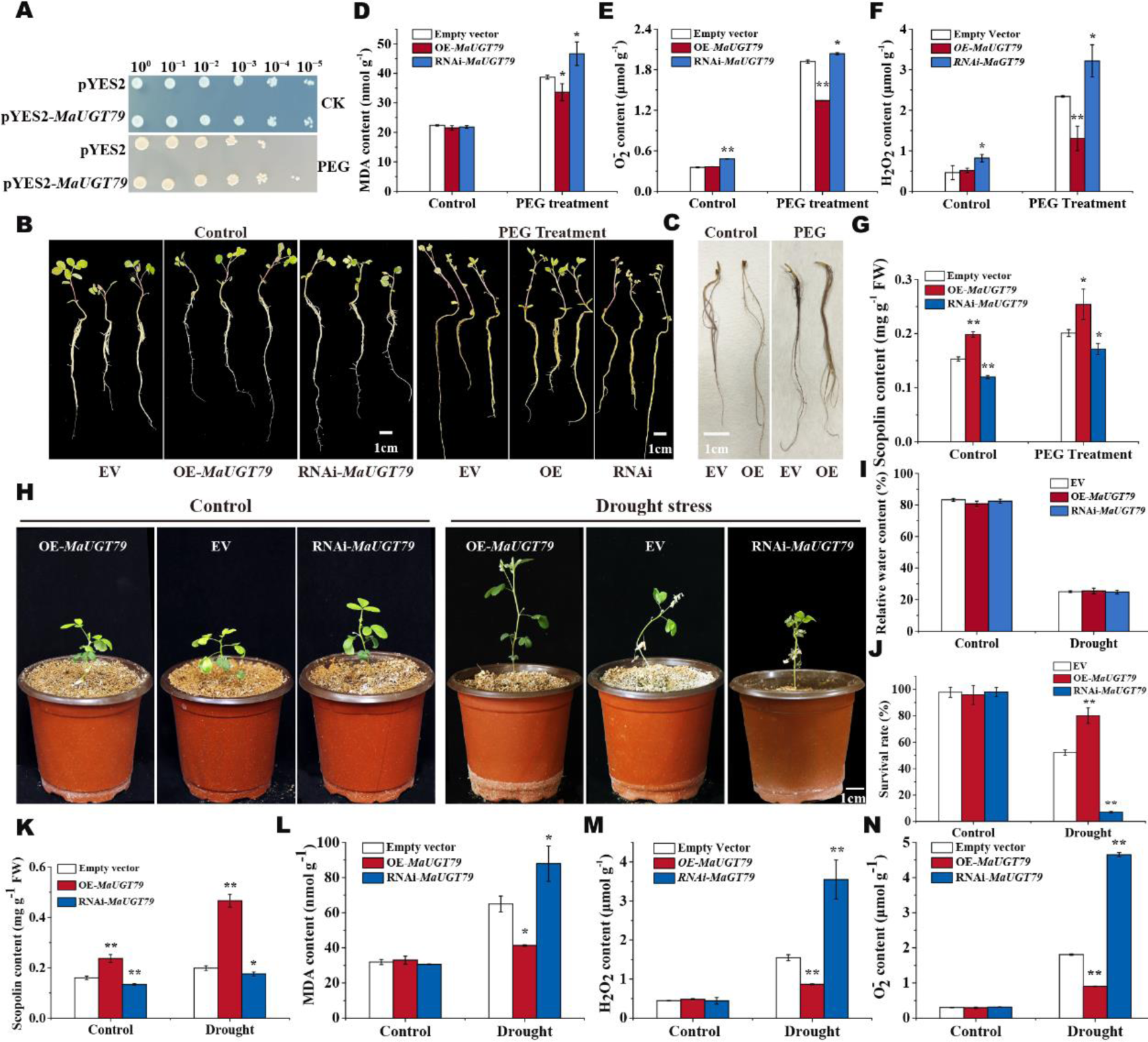
*MaUGT79* positively regulates drought tolerance in *M. albus* hairy roots. A) Drought stress tolerance analysis of *MaUGT79* in a yeast expression system compared with empty vector pYES2 (control) yeast. The two yeast cultures were independently grown in synthetic complete (SC)-Ura liquid medium containing 2% (*m/v*) galactose at 30 °C for 36 h up to A_600_=0.4. Then, the yeast was collected and adjusted with SC-Ura including 2% galactose and cultivated up to A_600_=1 for stress analysis. The same number of cells was resuspended in 30% PEG6000. Then, serial dilutions (10^0^, 10^-1^, 10^-2^, 10^-3^, 10^-4^, 10^-5^) were spotted onto SC-Ura agar plates and incubated at 30°C for 3 d. As a control, yeast with A_600_=1 without any stress was also spotted onto SC-Ura agar plates with the same dilutions as the treatments and grown at 30 °C for 3 d. B) Phenotypes of single seedling of overexpressing *MaUGT79* transgenic hairy roots (OE- *MaUGT79*), RNAi-*MaUGT79* transgenic hairy roots and transgenic hairy roots containing an empty vector (EV) grown under normal and 30% PEG6000 treatment for 3 days. C) Histochemical staining with NBT in hairy roots of EV and OE-*MaUGT79* under normal and 30% PEG6000 treatment for 3 days. (d-g) MDA content D), O2^-^ content E), H_2_O_2_ content F) and scopolin content G) in hairy roots of EV, OE- *MaUGT79* and RNAi-*MaUGT79* under normal and 30% PEG6000 treatment for 3 days. H) Phenotypes of single plant with OE-*MaUGT79*, RNAi-*MaUGT79* and EV transgenic hairy roots grown under normal and drought stress for 22 days. I) Relative water content of the plants with OE-*MaUGT79*, RNAi-*MaUGT79* and EV transgenic hairy roots under normal and drought stress at 22 days J), Survival rates of the plants with OE-*MaUGT79*, RNAi-*MaUGT79* and EV transgenic hairy roots grown under drought stress at 22 days, K-N) scopolin content K), MDA content L), O2^-^ content M), H_2_O_2_ content N) in hairy roots of EV, OE-*MaUGT79* and RNAi-*MaUGT79* grown under normal and drought stress for 22 days. The error bars indicate the SD values from at least three repetitions of each treatment. Asterisks indicate significant differences between EV, OE-*MaUGT79* and RNAi-*MaUGT79* under the same growth conditions. Significant differences were detected by Student’s t-test: *, *P*<0.05 or **, *P*<0.01.

### MaMYB4 activates *MaUGT79* expression through binding to its promoter

To understand the transcriptional regulation of *MaUGT79*, the upstream promoter regions (2.0 kb in size) of *MaUGT79* genes were analyzed for the prediction of potential *cis-*elements. We found that the promoter region contained many MYB binding sites (Figure 6A). Numerous studies have clearly demonstrated that MYB transcription factors function as key regulators of plant secondary metabolism (Chen *et al*., 2019). One of the very few works dealing with the regulation of scopolin production by transcription factors is about the AtMYB4 promotion of scopoletin production (Schenke *et al*., 2011). So, a phylogenetic tree comprising the sequences of amino acids of AtMYB4 (At4g38620) and MYBs of *M. albus* identified previously (Chen *et al*., 2021) showed that *Malbus0702723.1* was most closely related to AtMYB4, as they shared 82.35% amino acid sequence identity (Figure S6A). We thus designated this protein as MaMYB4. qRT-PCR analysis also showed that the expression of MaMYB4 was also induced by 30% PEG treatment (Figure S6B). The MaMYB4 transcript appeared mainly to be located in the nucleus (Figure S6D), which is consistent with its putative role as a transcription factor in the nucleus.

**Figure 6.**
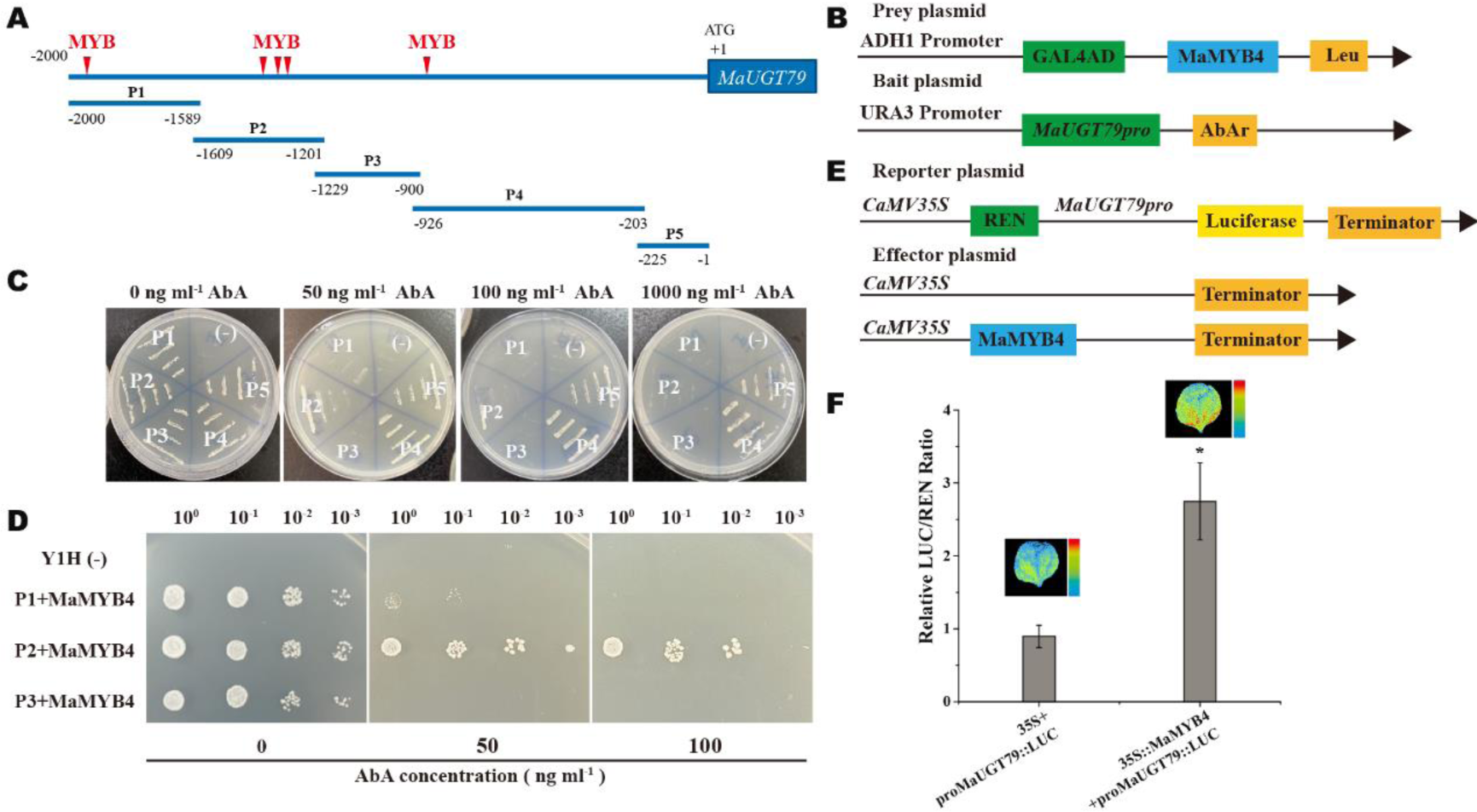
The promoter of *MaUGT79* is the direct target of MaMYB4 (MYB, myeloblastosis). A) Schematic diagram of the bait fragments (P1 to P5) used to construct the reporter vectors in the yeast one-hybrid assay. The red triangles indicate MBY biding sites. B) Schematic diagram of the prey plasmid and bait plasmid in yeast one-hybrid (Y1H) assay. The promoter fragment of *MaUGT79* was cloned into the pAbAi vector to generate the bait plasmid and the prey plasmid was generated by recombining the MaMYB4 gene into the pGADT7 vector. C) Transcriptional activation analysis of Y1H Gold [*Pro1/2/3/4/5/MaUGT79*-pAbAi]. D) Yeast one-hybrid assay. A pair of plasmids, pAbAi containing different fragments of the *MaUGT79* promoter and pGADT7 containing MaMYB4 were introduced into yeast strain Y1H gold and cultured on SD medium without Leu containing different concentrations of AbA at 30°C for 3 days. E) Schematic diagram of the reporter plasmid and effector plasmid. The promoter fragment of MaUGT79 was cloned into the pGreenII 0800-LUC vector to generate the reporter plasmid. The effector plasmid was generated by recombining the MaMYB4 gene into an overexpression vector (pBI 121). F) Dual-luciferase (LUC) assay in *N. benthamiana* leaves showing that MaMYB4 activates transcription of *MaUGT79* promoters. The leaves infiltrated with the empty vector and *MaUGT79pro::LUC* as a control. Representative photographs were taken (above), and LUC/Renilla Luciferase (REN) activity detection to verify that MaMYB4 activates the transcription of *MaUGT79* (below). The error bars indicate the SD values from the mean of at least five repetitions. Significant differences were detected by Student’s t- test: *, *P*<0.05.

In order to confirm whether *MaUGT79* is regulated by MaMYB4, we first conducted a Y1H assay. A 2000-bp DNA sequence upstream of the *MaUGT79* start codon was separated into five parts, P1 (-2,000 to -1,589), P2 (-1,609 to -1,201), P3 (-1,229 to - 900), P4 (-926 to -203), and P5 (-225 to -1, Figure 6A). We then integrated the P1, P2, P3, P4 and P5 sequences individually into the genomes of yeast cells. After introducing pGADT7-MaMYB4 into each of the respective yeast strains, we found that the P4 and P5 sequences of the *MaUGT79* promoter were not suitable for the Y1H system because 1000 ng/mL AbA was still unable to suppress the basal expression in the Y1H Gold harbouring *P4MaUGT79*-AbAi and *P5MaUGT79*-AbAi (Figure 6C), and then only one strain, carrying the P2 promoter, was able to grow on selective media (Figure 6D). This indicated that MaMYB4 binds to the P2 fragments of the *MaUGT79* promoter.

Subsequently, a dual-luciferase reporter assay was performed to further verify whether MaMYB4 activates the expression of *MaUGT79*. The *MaUGT79* promoter was used to drive the luciferase (LUC) gene as fusion reporters, with MaMYB4 overexpressed under the control of the CaMV *35S* promoter as an effector (Figure 6E). As shown in Figure 6f, the MaMYB4 and *MaUGT79* promoter co-transfected tobacco had a 2.7-fold higher relative LUC/REN ratio than the control, supporting the concept of an interaction between MaMYB4 and the *MaUGT79* promoter. Detection of LUC luminescence indicated that co-expression with the MaMYB4 transcription factor increased the expression of the *MaUGT79pro::LUC* reporters compared to the control lacking the *35Spro::MaMYB4* (Figure 6F). These results indicated that MaMYB4 can thus transcriptionally up-regulate *MaUGT79*.

### MaMYB4 positively regulates *MaUGT79*-mediated scopolin accumulation and drought tolerance

In order to explore the MaMYB4 expression profiles of *M. albus*, qRT-PCR was performed to assess transcript accumulation in different tissues. MaMYB4 was highly expressed in the leaves, which was positively correlated with the expression of *MaUGT79* (Figure S6B) and was induced by drought stress (Figure S6C). In order to gain further understanding of the regulatory roles of MaMYB4, overexpression and RNAi transgenic hairy roots were generated (Figure 7). qRT-PCR confirmed that the transgenic hairy roots accumulated high levels of MaMYB4 transcripts (2.9- to 5.3-fold; Figure 7A), which in turn increased the expression levels of *MaUGT79* by 8.9- to 20.4- fold (Figure 7B). Scopolin content increased by 1.66-, 1.54-, and 1.68-fold in comparison to the controls, respectively (Figure 7C). RNAi transgenic hairy roots exhibited low expression levels of MaMYB4 (0.43- to 0.58- fold), which in turn decreased the expression levels of *MaUGT79* by 0.29- to 0.47- fold. Scopolin content decreased by 0.91-, 0.78- and 0.92- fold compared with the controls, respectively. Interestingly, the transcription of *CCoAOMT1* and *F6’H1* genes involved in scopolin biosynthesis were obviously enhanced in OE-MaMYB4 lines and declined in RNAi lines (Figure 7D). Taken together, these findings support the assumption that MaMYB4 is a positive factor that modulates scopolin biosynthesis.

**Figure 7.**
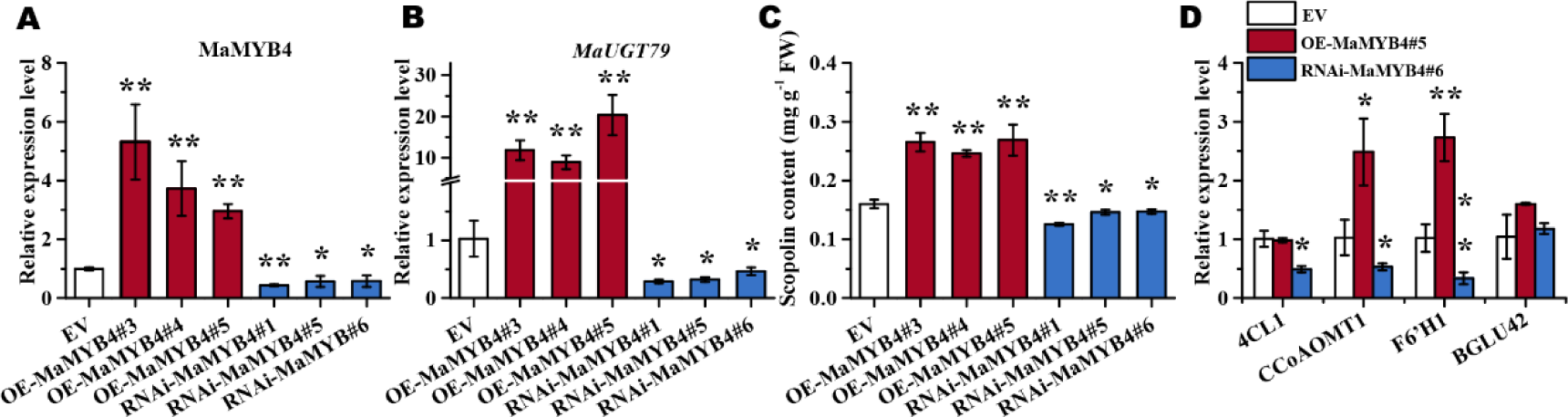
MaMYB4 positively regulates scopolin content in *M. albus* hairy roots. (MYB, myeloblastosis). A-B) Quantitative reverse-transcription (qRT)-PCR analysis of MaMYB4 A) and *MaUGT79* B) in control (EV), OE-MaMYB4 and RNAi-MaMYB4 transgenic hairy roots. C) Scopolin content in control (EV), OE-MaMYB4 and RNAi- MaMYB4 transgenic hairy roots. D) Gene expression levels of four scopolin biosynthesis genes (*4CL1*, *CCoAOMT1*, *F6’H1*, and *BGLU42*) in 21-d-old hairy roots. qRT-PCR was performed to detect gene expression levels. Data are normalized by β*- tubulin*. The error bars indicate the SD values from the mean of at least three repetitions. Significant differences were detected by Student’s t-test: *, *P*<0.05 or **, *P*<0.01.

Further, OE-MaMYB4 and RNAi-MaMYB4 transgenic hairy roots were selected to investigate whether MaMYB4 plays a role in drought tolerance, where transgenic hairy roots containing an empty vector were used as the control (EV). The EV, OE and RNAi transgenic hairy roots were subjected to 30% PEG6000 treatments for 3 days and water-deficit treatment for 22 days (the relative water content was 24.72%). Expectedly, the plants with OE-MaMYB4 transgenic hairy roots had significantly improved drought tolerance, while more severe wilting and necrosis of the leaves were observed in the plants with RNAi transgenic hairy roots compared with the EV (Figure 8A, H). The survival rates of the plants with MaMYB4-overexpressing transgenic hairy roots were 78.57-86.67%, whereas only 6.69% of RNAi-MaMYB4 transgenic hairy roots plants survived. Control plants had a 56.83% survival rate 22 days after drought stress was applied (Figure 5J), which was evidenced by the results of NBT and DAB staining (Figure 8B, C), and MDA, H_2_O_2_ and O_2-_ contents (Figure 8D-F, L-N). In addition, after 30% PEG6000 treatment, the expression level of *MaUGT79* in OE-MaMYB4 transgenic hairy roots was significantly up-regulated, while it was significantly inhibited in RNAi-MYB4 transgenic hairy roots (Figure 8G), indicating that MaMYB4 expression was up-regulated under drought stress and MaMYB4 activates *MaUGT79* expression. Accordingly, the content of scopolin increased significantly under drought stress in OE-MaMYB4 transgenic hairy roots (Figure 8K). Consistently, drought marker genes, such as *MaCOR47*, *MaRD29A*, *MaLEA3*, *MaP5CS1*, *MaRD29B* and *MaDREB2B*, exhibited significantly higher expression levels in OE-MaMYB4 hairy roots and lower expression levels in RNAi-MaMYB4 hairy roots upon exposure to 30% PEG6000 treatment (Figure S7). Taken together, these results indicated that MaMYB4 also plays a positive role in drought tolerance.

**Figure 8.**
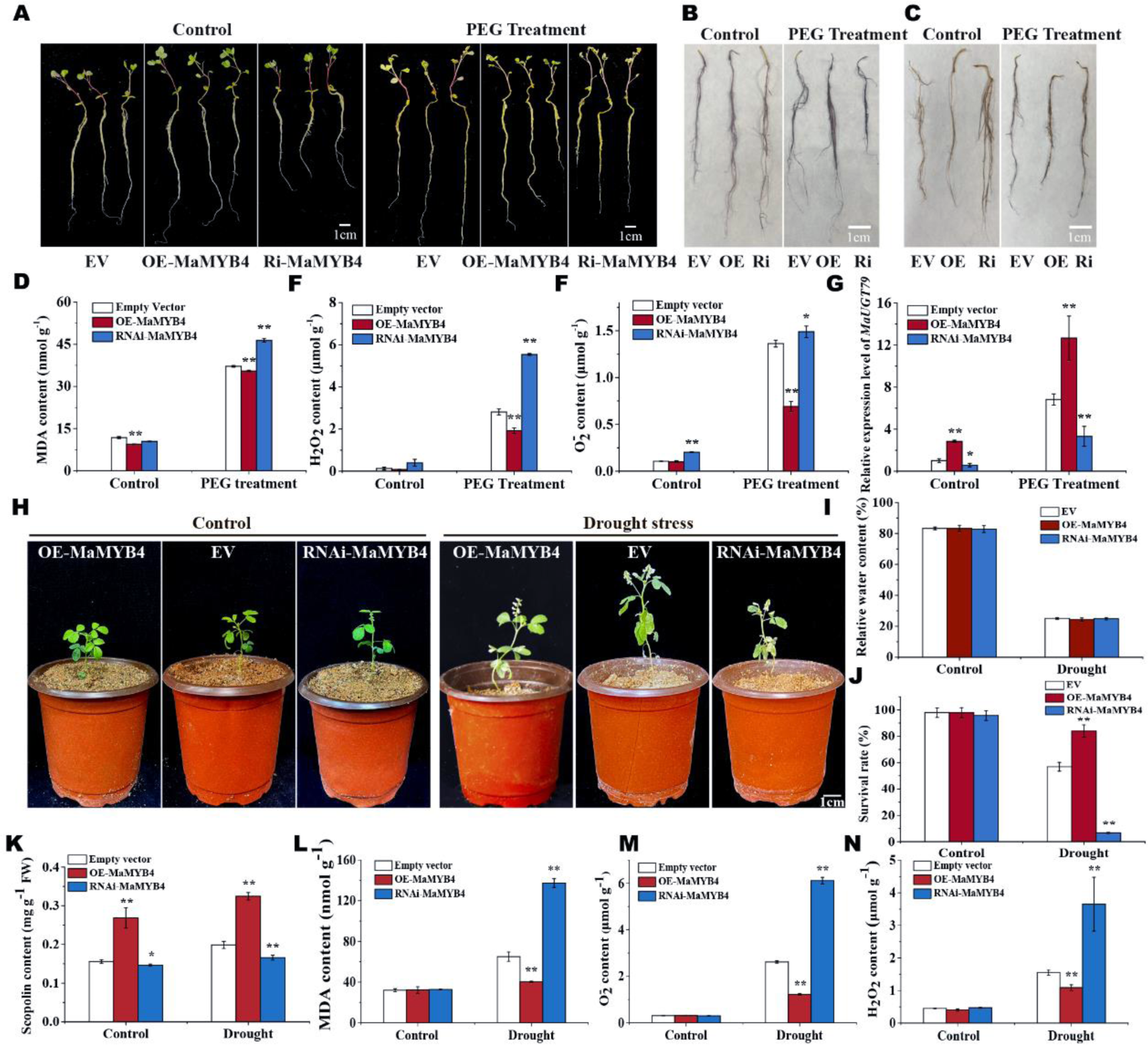
MaMYB4 positively regulates drought tolerance in *M. albus* hairy roots. A) Phenotypes of single seedling of overexpressing MaMYB4 transgenic hairy roots (OE- MaMYB4), RNAi-MaMYB4 transgenic hairy roots and transgenic hairy roots transferring an empty vector (EV) under normal and 30% PEG6000 treatment for 3 days. B-C) Histochemical staining with NBT B) and DAB C) in hairy roots of EV, OE- MaMYB4 and RNAi-MaMYB4 under normal and 30% PEG6000 treatment for 3 days. D-G) MDA content D), H_2_O_2_ content E), O2^-^ content F) and *MaUGT79* expression level G) in hairy roots of EV, OE-MaMYB4 and RNAi- MaMYB4 under normal and 30% PEG6000 treatment for 3 days. H) Phenotypes of single plants with OE-MaMYB4, RNAi- MaMYB4 and EV transgenic hairy roots under normal and drought stress for 22 days. I) Relative water content of plants with OE- MaMYB4, RNAi- MaMYB4 and EV transgenic hairy roots under normal and drought stress at 22 days J), Survival rates of plants with OE-MaMYB4, RNAi-MaMYB4 and EV transgenic hairy roots grown under drought stress at 22 days, K-N) scopolin content K), MDA content L), O2^-^ content M), H_2_O_2_ content N) in hairy roots of EV, OE-MaMYB4 and RNAi-MaMYB4 grown under normal and drought stress for 22 days. The error bars indicate the SD values from at least three repetitions of each treatment. Asterisks indicate significant differences between EV, OE-MaMYB4 and RNAi-MaMYB4 under the same growth conditions. Significant differences were detected by Student’s t-test: *, *P*<0.05 or **, *P*<0.01.

## Discussion

Using BSA based on 122,318 SNPs obtained from two NILs, JiMa46 and JiMa49 (Wu *et al*., 2022), we successfully identified a single polymorphic locus in a gene associated with scopolin biosynthesis that is located on chromosome 5, which we name as *MaUGT79.* To provide further insight into the genetic function and regulation mechanism of *MaUGT79*, we performed transgenic assays, substrate feeding assays and molecular biology experiments to demonstrate that *MaUGT79* functions as a positive regulator in scopolin biosynthesis of *M. albus* and enhances tolerance to drought stress. Our findings also highlight the regulatory function of a MYB transcription factor, MaMYB4, in scopolin biosynthesis and drought tolerance, and provide important insights into the regulatory mechanism underlying scopolin accumulation and drought tolerance in *M. albus*.

### The contribution of *MaUGT79* to drought stress tolerance is closely associated with scopolin accumulation

Previous studies have localized the global fluorescence of scopolin in leaf, stem, and root and found that coumarins move throughout the plant body via the xylem sap and it is a highly complex and dynamic process (Robe *et al*., 2021). Our result showed higher levels of scopolin in leaf, stem and root of *M. albus*. In Arabidopsis, BGLU42 was shown to be responsible for the deglycosylation of scopolin (Stringlis *et al*., 2018), but which is responsible for the glycosylation of scopolin is unkown (Robe *et al*., 2021).

In our study, *MaUGT79* encodes a UDP-glycosyltransferase that grouped in the same phylogenetic clade as AtUGT89B1 and AtUGT89A2, which belong to group B in our analysis (Figure 2B). AtUGT89A2 is a key factor that affects the differential accumulation of dihydroxybenzoic acid glycosides in arabidopsis (Chen & Li, 2017). A previous study showed that known coumarin UGTs, belonging to groups D (Langlois-Meurinne *et al*., 2005) and E (Huang, X-X *et al*., 2021), are more likely to be responsible for coumarin glycosylation modification. The group B protein, which is closely related to group D and E, in *M. albus* led us to propose *MaUGT79* as a major candidate gene for coumarin biosynthesis. Transgenic OE-*MaUGT79* in *M. albus* hairy roots indeed showed significantly increased scopolin accumulation (Figure 4). In Arabidopsis, scopolin accumulation in leaves was also reported in response to biotic and abiotic stresses (Doll *et al*., 2018). So we then closely examined the effects of overexpression of *MaUGT79* in response to drought stress. We first confirmed the positive role of scopolin in drought stress by reducing oxidative damage and promoting ROS scavenging (Figure S3). Then we found that the OE*-MaUGT79* lines displayed no obvious damage compared with the EV control under 30% PEG treatment and water-deficit treatment for 22 days (Figure 5B, H). In addition, OE-*MaUGT79* had significantly decreased MDA (Figure 5D, L), O2^-^(Figure 5E, N), and H_2_O_2_ (Figure 5F, M) contents, and increased scopolin content (Figure 5G, K) compared with the EV under 30% PEG6000 treatment and drought stress treatment. Further to this, the RNAi- *MaUGT79* transgenic lines showed an opposite trend (Figure 5D-F, L-N). We speculate that increased drought tolerance may be conferred by the correlated *MaUGT79*- mediated scopolin accumulation. Similar effects of coumarin-accumulation on increased abiotic stress have been reported in *Salvadora persica*, rice (*Oryza sativa*) and peanut (*Arachis hypogaea*) (Qin *et al*., 2019; Rangani *et al*., 2020; Patel *et al*., 2021). We conclude that the increased scopolin content in OE-*MaUGT79* hairy roots strengthens the ROS scavenging activity of the *M. albus* hairy roots and subsequently increases drought tolerance.

### Glycosylation of scopoletin promotes scopoletin biosynthesis via feedback activation of scopoletin biosynthesis genes

In our study, we observed that *MaUGT79* overexpression lines had greatly upregulated expression of critical genes encoding enzymes involved in scopoletin biosynthesis, including *4CL1*, *CCoAOMT1*, *F6’H1* and *BGLU42*, whereas the knock-down lines of *MaUGT79* had decreased transcription of these genes. These findings demonstrate that glycosylation of scopoletin accelerated the biosynthesis of the scopoletin, and that this is closely correlated with the upregulation of the biosynthesis genes. A previous study showed that overexpression of *TOGT*, a scopoletin glucosyltransferase, in tobacco results in both scopoletin and scopolin over-accumulation as compared to wild-type (Gachon *et al*., 2004), suggesting that up-regulation of glycosylating activity toward a specific substrate does not necessarily result in lower accumulation of the corresponding aglycone form. So, we believe that as the plant cells continuously consume aglycones upon constitutive expression of *MaUGT79*, they require more substrate, which in turn stimulates the expression of the upstream enzyme encoding genes and accelerated biosynthesis of scopoletin.

### MaMYB4 is linked to scopolin metabolism via *MaUGT79* in regulating drought stress adaption

Transcription factors (TFs) are a group of regulators that play crucial roles in many plant biological and developmental processes by regulating gene expression at the transcriptional level through recognition of specific DNA sequences in promoters (Mitsuda & Ohme-Takagi, 2009). Although the biosynthesis of scopolin is most responsive to MaUGT79 activity in *M. albus*, knowledge of the mechanisms involved in the regulation of *MaUGT79* transcription is fairly limited. The TF MYB15 is proposed to regulate the basal synthesis of scopoletin (Chezem *et al*., 2017). MYB72, which tightly regulates *F6’H1* expression, is also involved in scopolin accumulation (Stringlis *et al*., 2018). In this study we identified a MYB TF, MaMYB4, whose expression closely correlates with *MaUGT79* gene expression (Figure S6), with overexpression leading to an increase in scopolin (Figure 7C). In grapes (*Vitis vinifera*), VvMYB4b, VvMYB4a, and VvMYB4-like were associated with reduced proanthocyanidin and anthocyanin accumulation and, down-regulation of structural and regulatory genes of the flavonoid biosynthesis pathway (Cavallini *et al*., 2015; Ricardo Perez-Diaz *et al*., 2016). In *A. thaliana*, AtMYB4 and AtMYB7 are two members in subgroup 4 of the R2R3-MYB transcription factors, overexpression of AtMYB4 reduces the expression of AtMYB7, and the lack of AtMYB7 results in an increase in the expression of the early phenylpropanoid genes *C4H*, *4CL*, and *UGT* (Fornale *et al*., 2014). These findings suggest that a variety of MYB4 sequences in different plant species are involved in various metabolic mechanisms, and thus a new function in regulating scopolin biosynthesis is presented in this study. We identified that MaMYB4 could directly control the expression of *MaUGT79*, a glycosyltransferase involved in modulating scopolin biosynthesis. Over-expression of MaMYB4 significantly enhanced the expression of *MaUGT79*, the content of scopolin and drought tolerance, and these levels were reduced when MaMYB4 was down-regulated via RNA-interference (Figure 8). Yeast one-hybrid (Y1H) and Dual-luciferase (LUC) assays showed that MaMYB4 acts by binding to the promoter of *MaUGT79* and activates *MaUGT79* transcription (Figure 6). These results link MaMYB4 to the scopolin biosynthetic pathway in improving drought stress tolerance through activating the expression of *MaUGT79*. Here, we add new knowledge about upstream regulatory factors of *MaUGT79*, which show developmental-based expression to stimulate scopolin accumulation and drought tolerance in *M. albus*.

In summary, we show that *MaUGT79* over-expression in *M. albus* hairy roots resulted in increased scoplin accumulation, leading to enhanced drought tolerance. Furthermore, we show that MaMYB4 over-expression promotes the deposition of scopolin by directly regulating the expression of *MaUGT79*. Under drought conditions, MaMYB4 expression is higher and MaMYB4 directly activates the expression of *MaUGT79*. MaUGT79 catalyses the conversion of scopoletin to scopolin, leading to an increase in scopolin content. The resulting accumulation of scopolin contributes to drought tolerance by increasing ROS scavenging capacity (Figure 9).

**Figure 9.**
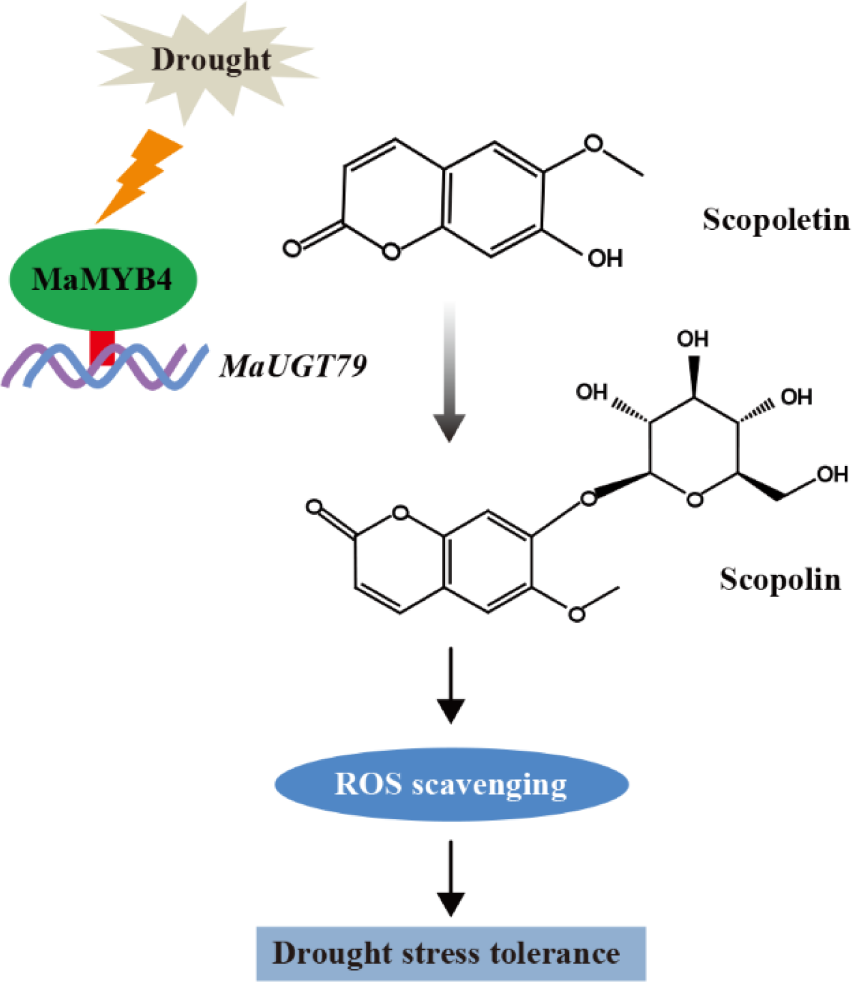
A proposed mechanistic model for the drought-dependent regulation of scopolin biosynthetic *MaUGT79* gene expression, regulated by MaMYB4 in *M. albus*. Under drought conditions, MaMYB4 is up-regulated and activates the expression of *MaUGT79*. MaUGT79 catalyses the conversion of scopoletin to scopolin, which results in an increase in scopolin content. The accumulation of scopolin leads to drought tolerance by increasing ROS scavenging.

## Materials and methods

### Plant materials and growth conditions

*M. albus* plants were grown in an illuminated incubator under controlled conditions (16 h/8 h day/night cycle at 25℃, with a relative humidity of 40%). Roots, stems, leaves, and flowers at the flowering stage were collected, frozen in liquid nitrogen and then stored at –80℃ until use. *Nicotiana benthamiana* plants used in this study were grown in pots in a growth chamber under a 16 h photoperiod with a 20°C: 25°C, night: day temperature.

### Bulked-segregant analysis sequencing (BSA-seq)

BSA-seq analysis was performed on the NILs to identify the target gene in this study. The two bulks consisted of the JiMa46 pool with an extremely low scopolin content phenotype and the JiMa49 pool with an extremely high scopolin content phenotype, and they were constructed by mixing an equal amount of DNA extracted from 30 individuals of JiMa46 and JiMa49 plants, respectively. SNP discovery was performed as described in a previous study (Wu *et al*., 2022).

### Gene cloning and sequence analysis

A 1343 bp and 1392 bp CDS of *MaUGT79* from JiMa46 and JiMa49, respectively, was amplified based on the genome data using Phanta Max Super-Fidelity DNA Polymerase (Vazyme Biotech Co.,Ltd, Nanjing). Gene specific primers (Table S1) were designed using DNAMAN software (Wang, 2015). The CDS was confirmed by sequencing. For phylogenetic analysis, the deduced amino acid sequences of MaUGT79 were aligned with previously characterized UGTs using multiple sequence comparison by ClustalX (Larkin *et al*., 2007) . The phylogenetic tree was constructed based on the alignments using MEGA 7.0 (Kumar *et al*., 2016) with the neighbour-joining (NJ) method, and bootstrap tests were performed using 1,000 replicates to support statistical reliability. A 2.0 kb sequence of the promoters of *MaUGT79* was cloned and used to identify the *cis*-acting elements in the promoter regions through the PlantCARE website (http://bioinformatics.psb.ugent.be/webtools/plantcare/html/) (Lescot *et al*., 2002).

### Subcellular localization assay

To investigate the subcellular localization of MaUGT79, we constructed a recombinant MaUGT79 tagged at the N-terminus with red fluorescent protein (RFP), and the CDS of *MaUGT79* was inserted between the *Xba*I and *Bam*HI sites of the binary vector pBI121 DsRed2 (Wu *et al*., 2022) with expression driven by the CaMV *35S* promoter. The specific primers are listed in Table S1. The leaves of 6-week-old *N. benthamiana* seedlings were selected for a transient overexpression experiment, *Agrobacterium tumefaciens* strain GV3101 carrying *35S:MaUGT79*-RFP and the control vector with RFP alone were transformed into the leaves according to a previous report (Liu, Xiaoying *et al*., 2022). The cytoplasm marker pBI-NLS-CFP was co-transformed into the *N. benthamiana* leaves to exhibit the location of the cytoplasm. The system was subsequently cultured in the dark for one day and then in daylight for another two days. A laser scanning confocal microscope (Olympus FV3000, Japan) was used to observe the image of transformed leaves.

### Heterologous protein expression and substrate feeding

The *MaUGT79* coding sequence was inserted between the *Bam*HI and *Sac*I sites of the pET32a vector (Duan *et al*., 2021) using the ClonExpress^®^ MultiS One Step Cloning Kit (Vazyme Biotech Co.,Ltd, Nanjing) following the protocol provided. Primers are shown in Table S1. The construct was then transformed into *Escherichia coli* strain BL21 (DE3) (TransGen Biotech, Beijing). After incubation at 37°C in Luria– Bertani (LB) liquid medium containing ampicillin (100 μg ml^-1^) for 24 h, the culture was diluted and grown until the optical density at 600nm (OD600) of the cultured cells reached 0.6-0.8. After adding 0.1 mM isopropyl-β-D-thiogalactopyranoside (IPTG) the culture was incubated at 16°C and 180 rpm for 20 h to induce expression of the recombinant protein (Sun *et al*., 2017). The protein was then purified using *Proteinlso^®^* Ni-NTA Resin (TransGen Biotech, Beijing) and analyzed on a 12% SDS-PAGE gel.

The BL21 (DE3) cells harboring the pET32a-*MaUGT79* vector and empty vector were cultured in the same conditions for *E. coli* expression. Afterwards, IPTG (final concentration of 0.1 mM) was added and incubated for 20 h, then the substrate *o*- coumaric acid, esculetin, umbelliferone, scopoletin (final concentration of 0.1 mM) and 500 mM UDP-Glucose (98%, Innochem Co., Ltd) were added, and the culture was incubated for 24 h. After that, the reaction system was extracted with an equal amount of ethyl acetate. The recovered ethyl acetate extract was decompressed and dried, then dissolved in MeOH for high-performance liquid chromatography (HPLC) analysis (Yang *et al*., 2016). The detection wavelength of *o*-coumaric acid and umbelliferone was 320 nm, the detection wavelength of esculetin was 350 nm, and the detection wavelength of scopoletin and scopolin was 346 nm.

### Validation of Heterologous Expression in Yeast

To produce the pYES2-*MaUGT79* construct, the full-length coding sequence of *MaUGT79* was amplified from JiMa49 and inserted between the *Bam*HI and *Xba*I sites of the pYES2 (Duan *et al*., 2021) expression vector using a ClonExpress^®^ MultiS One Step Cloning Kit following the manufacturer’s protocol with the specific primers listed in Supplementary Table S1. Following confirmation of the cloned sequence, the recombinant pYES2-*MaUGT79* plasmid and empty pYES2 plasmid were transformed into *Saccharomyces cerevisiae* strain INVSc1 (Duan *et al*., 2021). The two yeast cultures were independently grown in synthetic complete (SC)-Ura liquid medium containing 2% (*m*/*v*) galactose at 30°C for 36 h up to an OD600 of 0.4. Then, the yeasts were harvested and adjusted with SC-Ura including 2% galactose and cultivated up to an OD600 of 1.0 for stress analysis. The same amount of cells were resuspended in 30% PEG 6000 (Zhang *et al*., 2021). The treated yeast liquid was diluted 1:10 and cultured on SC-U/2% (*w*/*v*) glucose agar plates for 2–3 days to observe colony growth, and photos were taken to record the expression of the binding protein.

### Yeast one-hybrid assay

For Matchmaker Gold yeast one-hybrid system (Clontech, Mountain View, CA, USA), the MaMYB4 CDS was fused to the GAL4 transcription factor activation domain (GAL4AD) in the pGADT7 vector (Zheng *et al*., 2021) to generate the prey vector (pGADT7-MaMYB4). While the various promoter fragments of *MaUGT79* (2.0-kb) were inserted into the pAbAi vector (Zheng *et al*., 2021) to construct the baits (*pro1/2/3/4/5MaUGT79*-*AbAi*). These *Bst*BI-cut bait constructs were integrated separately into the genome of Y1HGold (Zheng *et al*., 2021) to generate five bait reporter strains. The minimal inhibitory concentrations of abscisic acid (AbA) were determined for the baits using SD/–Ura agar plates containing 0–1000 ng ml^-1^ AbA. After selecting the transformants on SD/–Ura plates, the pGADT7-MaMYB4 construct was introduced into the bait reporter strains, with a blank pGADT7 plasmid serving as a negative control. Positive transformants were selected on SD/–Leu medium supplemented with an appropriate concentration of AbA and cultured at 30°C for 3 d (Zheng *et al*., 2021).

### Dual-luciferase assay

For the dual-luciferase (Dual-LUC) assay, the *MaUGT79* promoter (-1609∼-1 bp) was ligated into pGreenII-0080-LUC (Zheng *et al*., 2021) to generate the reporter construct *proMaUGT79:LUC*. The MaMYB4 CDS was ligated into the binary vector pBI121 to generate the effector construct *35S:MaMYB4*. The effector and reporter constructs were transformed into *A. tumefaciens* strain GV3101 harbouring the pSoup helper vector (Nguyen *et al*., 2021), respectively, which were further co-infected into 6-week-old *N. benthamiana* leaves. The leaves were infiltrated with the pBI121 effector construct and the *proMaUG*T79*:LUC* as a control (Zheng *et al*., 2021). The injected tobacco plants were kept in the dark for 12 hours and then 2 days in normal light conditions. A Fluorescence Chemiluminescence Imaging System (FX6.EDGE Spectra; VILBER, France) was used to capture the LUC image. The promoter activities were determined by measuring Firefly Luciferase to Renilla Luciferase (LUC/REN) ratios using the Dual Luciferase Reporter Gene Assay Kit (RG027, Beyotime, Shanghai, China) with a Multimode Reader (Varioskan LUX, Thermo Fisher, Finland). Five biological replications were measured for each sample.

### Vector construction and *Agrobacterium rhizogenes*-mediated transformation system of *M. albus*

To produce the *MaUGT79* and MaMYB4 overexpression and RNAi expression constructs, the full-length cDNAs of *MaUGT79* and MaMYB4 were cloned into the binary vectors pBI121 and pK7GWIWG2 (II) RR, using the ClonExpress^®^ MultiS One Step Cloning Kit and Gateway LR Clonase Enzyme Mix (Invitrogen), respectively. The constructs were introduced into *A. rhizogenes* strain K599 by the electroporation method (Wang *et al*., 2021). Transgenic hairy roots were obtained according to a previous report (Wang *et al*., 2021). The transgenic hairy root lines containing the overexpression empty vector (EV) were used as a control. All transgenic lines were tested by PCR and RFP visualization to identify positive lines. The transgenic hairy roots were then transferred to Murashige & Skoog medium (with 100 mg ml^-1^ cefotaxime) an maintained in the dark, at 22°C, for two months. Hairy roots were then harvested for determination of scopolin content (Figure S2a).

### Drought stress and exogenous scopolin treatments and tolerance evaluation

In order to investigate the changes in *MaUGT79* expression under drought stress, six-week-old plants were treated with 30% PEG6000. The leaf samples for qRT-PCR analysis were harvested at 0, 3, and 24 h after treatment. All samples were immediately frozen in liquid nitrogen and stored at -80°C. Three replicates were performed for each sample. For *M. albus* hairy root drought experiments, the 2-month-old EV, OE-*MaUGT79*/MaMYB4 and RNAi-*MaUGT79*/MaMYB4 transgenic hairy roots lines were treated with 30% PEG6000 for three days. And the EV and RNAi*-MaUGT79* lines were also treated with 30% PEG6000+100 μM scopolin for three days. Moreover, after 18 days growth of the plants with transgenic hairy roots, the seedlings from each line were carefully transferred to flowerpots containing vermiculite and sand (*v*/*v* = 1:1) for 30 days of growth and then the seedlings were used in the phenotyping experiment. The plants grown under water-replete conditions were watered twice per week with 1/2 Hoagland nutrient solution. For drought tolerance comparisons, water was withheld from *MaUGT79*/MaMYB4-overexpressing, *MaUGT79*/MaMYB4-silencing and control lines as in Zhang *et al*. (2012) for 22 days until there were distinguishable differences between control lines and *MaUGT79*/MaMYB4-overexpressing or *MaUGT79*/MaMYB4-silencing lines. At least 30 plants per independent line were evaluated in each treatment, and all treatments were repeated three times.

### RNA extraction and gene expression analysis

Total RNA was extracted from leaves, stems and roots at the flowering stage and from hairy roots of *M. albus* using the TransZol reagent (TransGen Biotech, Beijing). First strand cDNA was obtained using the Hifair^®^ Ⅲ 1st Strand cDNA Synthesis SuperMix for qPCR (gDNA digester plus) by oligo(dT) primer (Yeasen biotech Co., Ltd., Shanghai). Quantitative RT-PCR was performed using Hieff^®^ qPCR SYBR^®^ Green Master Mix (No Rox) (Yeasen biotech Co., Ltd., Shanghai) on a CFX96 Real-Time PCR Detection System (Bio-Rad, Los Angeles, CA, USA). β*-tubulin* was used as a housekeeping reference gene. The expression levels were calculated relative to the reference and determined using the 2^-ΔΔCT^ method (Zong *et al*., 2021). There were three biological replicates for all analyses. Primers used for qRT-PCR are listed in Table S1.

### Scopolin extraction and quantification

For scopolin extraction, ambient temperature-dried samples derived from fresh samples were ground and passed through a sieve with an aperture size of 0.45 mm and extracted with an ethanol/water mixture (80: 20, *v/v*). For tissue extraction, 50 ml of solvent g^−1^ of dry weight was used. For hairy roots extraction, 5 ml of 80% ethanol per 100 mg was added to the frozen material (Doll *et al*., 2018). After shaking for 10 min, ultrasonic extraction was performed at room temperature for 60 min. The ratio between the weight of the fresh samples and the volume of the extraction solution was the same for all samples in a given experiment. The extracts were filtered through 0.45 µm filters for high performance liquid chromatography (HPLC) analysis.

HPLC separation was performed on an Agilent 1100 HPLC system using a 5 µm C18 column (4.6 mm *×* 150 mm, Agilent-XDB), maintained at 30°C, with water (containing 0.1% phosphorous acid) and acetonitrile as the mobile phase. The flow rate of the mobile phase was set at 1 ml min^-1^ over 20 min. For the quantification of scopolin, the calibration was performed with an eight-point calibration curve made using commercial sources of scopolin and scopoletin (Chengdu PureChem-Standard Co., Ltd) (Figure S7). Chemicals used in this study were of analytical or HPLC grade.

### Measurement of physiological and histochemical staining

For physiological analyses, *M. albus* plants were treated with 30% PEG6000 for 3 days. Histochemical staining of O_2-_ was conducted by the BCIP/NBT Chromogen Kit (Solarbio, Beijing, China). The MDA, O2^-^ and H_2_O_2_ content were measured using the detection kit (Solarbio, Beijing, China) according to the manufacturer’s instructions, respectively (Huang, X *et al*., 2021). Histochemical staining of H_2_O_2_ was conducted using the DAB Chromogen Kit (Solarbio, Beijing, China).

### Data Availability statement

The genomic data of *M. ablus* are openly available in NCBI (NCBI BioProject ID PRJNA674670).

## Acknowledgments

This work was financially supported by the National Natural Science Foundation of China (32061143035, 32271752), Inner Mongolia Seed Industry Science and Technology Innovation Major Demonstration Project (2022JBGS0040), the Fundamental Research Funds for the Central Universities (lzujbky-2022-ey17).

## Author contributions

JZ, ZD, FW and QY designed the experiment and conception; ZD, YW, SW, PZ and CZ performed experiments; ZD, FW, QY, SW, YW, CZ and JZ analyzed data. ZD, QY and SW wrote the manuscript. JZ and CSJ revised the manuscript. All authors read and approved the manuscript.

## Supplemental data

**Supplemental Table S1** List of primers used in the present study.

Supplemental Figure S1. **S**DS-PAGE analysis of MaUGT79 heterologously expressed in *E. coli*. Expression of the protein in *E. coli* BL21(DE3) was induced by IPTG for 20 h at 16℃. M, protein molecular size markers; Lane 1, empty vector; Lane 2, induction of MaUGT79; Lane 3, purified MaUGT79 protein.

Supplemental Figure S2. Phenotype and identification of transgenic hairy roots. A) Phenotype of transgenic hairy roots. B) Genomic DNA (gDNA) PCR identification of *35S:MaUGT79* in transgenic *M. albus* hairy roots using 35S-F/RED-R primers shown in supplementary table. C) RFP signals in *MaUGT79*-overexpressing hairy roots of *M. albus*.

Supplemental Figure S3. A) HPLC profiling of scopolin and scopoletin in control (EV), overexpression lines (OE-1 and OE-5), and RNAi lines (RNAi-1 and RNAi-6) under 30% PEG6000 treatment and applied with exogenous scopolin after 30% PEG6000 treatment. Authentic scopolin and scopoletin were used as standards. B-C) MDA content B) and O2^-^ content C) in hairy roots of EV and RNAi-*MaUGT79* under control (CK), 30% PEG6000 treatment and 30% PEG6000+scopolin treatment for 3 days. The error bars indicate the SD values from the mean of at least three repetitions of each treatment (*, *P*< 0.05 or **, *P* < 0.01).

Supplemental Figure S4. Expression pattern of *MaUGT79* under 30% PEG6000 treatment using qRT-PCR. The values shown are the means ± standard deviation of three replicates. β*-tubulin* was used as a normalization control for qRT-PCR. Red lines indicate the expression values (FPKM) from RNA-seq data. CK represents the control.

Supplemental Figure S5. Expression levels of drought marker genes including *MaCOR47* A), *MaRD29A1* B), *MaLEA3* C), *MaP5CS1* D), *MaRD29B* E) and *MaDREB2B* F) in hairy roots of EV and two OE-*MaUGT79* lines (OE-*MaUGT79*#1 and OE-*MaUGT79*#5) under control and 30% PEG6000 treatment for 3 days. β*-tubulin* was used as a control for qRT-PCR. The error bars indicate the SD values from the mean of at least three repeats of each treatment. Asterisks indicate significant differences (*, *P*< 0.05 or **, *P*< 0.01) between EV and OE-MaUGT79 under the same growth conditions based on Student’s *t*-test.

Supplemental Figure S6. Correlation and phylogenetic analysis of MaMBY4. A) Phylogenetic analysis of the amino acid sequences of AtMYB4 (At4g38620) and MYBs of *M. albus*. Gene ID for labels can be found in supplementary table. B) The correlation between the expression of MaMYB4 and *MaUGT79* in different tissues of *M. albus* at the flowering stage. Error bars represent± SD (n=3). C) Expression pattern of MaMYB4 under 30% PEG6000 treatment using qRT-PCR. The values shown are the means±standard deviation of three replicates. β*-tubulin* was used as the reference gene. D) The subcellular localization of MaMYB4. MaMYB4 was fused to RFP into *N. benthamiana* leaves. The fluorescence was observed under a confocal laser scanning microscope. The cyan signal of NLS (nuclear localization signal)-CFP shows the location of the nuclear marker. Scale bars indicate 50 µm.

Supplemental Figure S7. Expression levels of drought marker genes including *MaCOR47* A), *MaRD29A1* B), *MaLEA3* C), *MaP5CS1* D), *MaRD29B* E) and *MaDREB2B* F) in hairy roots of EV, OE-MaMYB4 and RNAi-MaMYB4 under control and 30% PEG6000 treatment when grown for 3 days. β*-tubulin* was used as normalization controls for qRT-PCR. The error bars indicate the SD values from at least three repeats of each treatment. Asterisks indicate significant differences (*, *P*< 0.05 or **, *P*< 0.01) between EV, OE-MaMYB4 and RNAi-MaMYB4 under the same growth conditions based on Student’s *t*-test.

Supplemental Figure S8. The calibration curve of scopolin.

